# In silicon emergence of an autonomous artificial metabolic system

**DOI:** 10.1101/865808

**Authors:** Shijian Chen

**Affiliations:** School of computer and information technology, Shanxi University, China

## Abstract

In this work, we establish and evolve an artificial metabolic system in silicon to shed light on how the metabolic mechanism emerged. This system is composed of two subsystems: the artificial genome subsystem (AGS) and the artificial metabolite subsystem (AMS). The whole system is designed to be capable of being autonomous: the dynamics of AGS is capable of situating itself to the dynamics of AMS to provide it with enzymes in the right time and quantity; the dynamics of AMS is capable of implementing the metabolic function and harvest energy so as to pay back the energy consumption of AGS. This kind of autonomous state requires an intricate structure of the AGS. So it is almost impossible to be predetermined manually. With the help of an evolutionary computational method that has a hierarchical mutational structure, the artificial metabolic system with this kind of autonomous state eventually emerged in silicon. We find that ATP and ADP molecules have an important role in making the state of the system autonomous. We also find that the emerged structure of AGS ensemble existing biological structures in the natural cells.

## 1. Introduction

Metabolic process is an intensively studied area. It is a complex process including many interacting metabolites. Take glycolysis and the citric cycle for instance, dozens of metabolites take part in the process, which forms a system called metabolic pathway. They have chain or cyclic structures etc. On the other hand, behind the metabolic path way lies another system which maintains and controls the process. This system absorbs the energy and material from the metabolic pathway and constantly gives back enzymes or control signals to maintain the operation of the process. The system is called the gene regulatory network. A lot of research has also been done in this area, and it is revealed that GRN also have complex structures. There are motifs such as feedback or feed forward motifs [1] or modular structures [2].

There have been some works using computational method to investigate the origin of the mechanism of metabolism[9]. In [3], they evolve their system in silico and make a metabolic process of 33 metabolites to emerge. But in their work, enzymes are available whenever needed without considering how GRN regulates this process. In [4] The author uses 6 segment of fixed length binary string to represent a gene. The dynamics of this article is extended to concentration profiles, oscillating dynamics and signal responses. This article did not incorporate the dynamics of a relatively independent biochemical process that need the protein system to regulate. These works simplified these two processes in certain level. And as they usually only focus on one subsystem, the intimate interaction between them are lost.

In our opinion, GRN and metabolic pathway is likely to have co-evolved. So separating them may lose essential clues. So we make an artificial metabolic system which contains both of them. The consequence is that the system has the ability to regulate its own biochemical process, which makes this system to be autonomous.

We use an artificial genome model which mimics the structure of natural DNA molecule. It only defines the basic rules that inheritance information is stored, mutated and dynamically expressed. The metabolic reaction model well recovers the essence of the biochemical process in the cell, that is, all information or energy transmission has to be done by the collision of molecules. The gene regulation process and the metabolic process are all dynamical and self-sufficient based on simple molecular interactions, which preserves their capacity to intimately interact with each other. Having these two subsystems to run in parallel, we have the hope to peek in to the secret of how they co-evolve into a system with meaningful structures.

The article is organized as follows. First we introduce the artificial genome model. Second is the mutational mode of artificial genome model. Third is catabolic reaction model. The last part is the construction and testing of the whole integrated system. Then we develop the hierarchical evolutionary algorithm to evolve this system. Eventually we analyze the final evolved transcriptional regulatory network and the dynamical process.

## 1. Artificial genome model and realization of concurrency

### 1.1 Artificial genome model

The artificial genome is represented as a bit string, displayed in Fig 1. It is based on the model used in [5]. The detailed description of it can be seen in the affiliation part.

**Fig 1.**
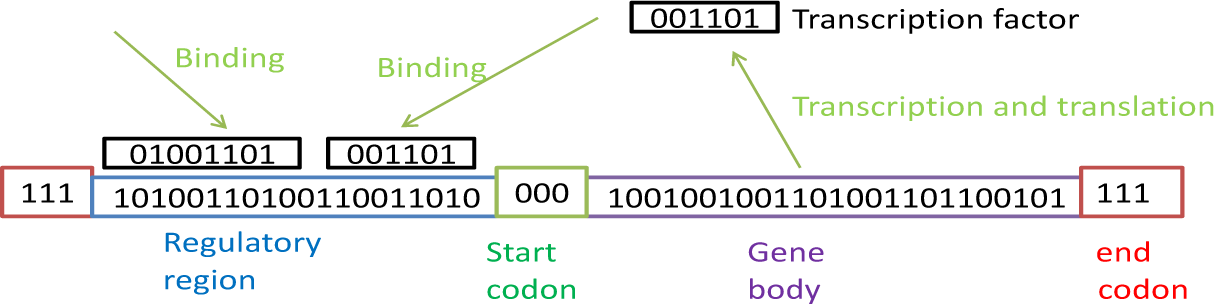
Illustration of artificial genome.

We allow multiple transcription factors to bind to the transcription regulatory zone as illustrated in Fig 1. To get a single control signal out of multiple transcription factors, we concatenate all transcription factors in the same zone together. Then we use a hash function to operate on this sequence and decide whether the control signal is on or off due to the hash value as illustrated in Fig 2. This way, the adding or removing of any single or combination of transcription factors has the potential to turn on or off the gene transcription, which makes the regulation mode much flexible.

**Fig 2.**
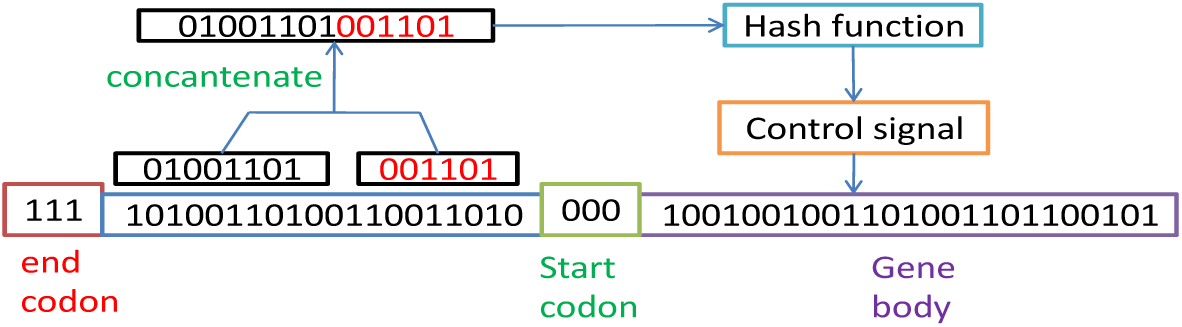
Control signal generation.

The basic data structure of the genome and the regulation mode has been established. Next we need to establish the rule which makes the system operates dynamically. In a nature cell, a transcription factor diffuses in the cytoplasm and searches alone the DNA molecule for its binding site by complex biochemical interactions [6]. To our knowledge, this process is very efficient, so we simplify it to be an instant event soon after the TF is made. If there are multiple binding sites available, we just randomly select one of them. We are satisfied with this level of proximity.

The degradation of proteins is another important factor for the process, but is easy to be ignored. When degradation happens is regulated by N-degron or C-degron pathways as reported in[7]. They determine both the life times of TFs and enzymes, which then influence the dynamics of the whole system. For simplicity, we regard the life time of each protein to be constant. We use the last 11 digits of the protein sequence to determine its life time. What’s more we make this parameter to be specific to each protein and evolvable. This will empower the artificial genome with great evolutionary potential.

### 1.3 Realization of concurrency

As we know, all the transcription and regulation process in the a natural cell happens concurrently. Parallel computation is a natural choice. But the interactions of genes on the DNA are intricate, making the implementation of parallel computation cumbersome. So we chose to use a serial computation to make the concurrency happen.

For each gene which has been activated or TF which binds to the genome, there is a positive integer number which is called a timer corresponds to it. The binary string of the genome is scanned repeatedly for genes or TFs with timers. A timer which is scanned will be reduced by 1 unit. An event on the genome is triggered by a timer counted down from 1 to 0. Once triggered, execute the event and precede the scan process. The timer of a gene triggers the binding of a TF it has made to the genome and set the timer of its controlling gene, and that of the TF triggers its degradation and getting off from the genome. In this way, after one round of scan, all events on the DNA happen. If we regard that the events in this round happens at the same time, then the concurrency is accomplished. One round of scan actually defines how time elapses for one unit on the genome. Mathematical description of this method is in the method part.

## 3. Catabolic function model

Another part of our system is the artificial metabolic subsystem. This subsystem directly absorbs sugar from the environment and supplies the energy consumption of the whole system. On the other hand, the operation of this subsystem relies on the constant and appropriate supply of enzymes from the artificial genome subsystem. The metabolic pathway is not predetermined. We just define what kind of molecules there are and the way they interact. The dynamical model mimics the biochemical reaction in a stirred tank, rather than a predetermined ODE model or other discrete models.

The modeling of biochemical reaction relies on the change of free energy of reactants catalyzed by enzymes. So we first define how energy is computed by the arrangement of its string. We consider each digit and its neighbor has an artificial chemical bond which has potential energy. 11 and 00 all have energy of 1, while 10 and 01 all have energy of -1. The energy of a sugar string is the sum of all the energy of these digit pairs in the string. An example is illustrated in Fig 3.

**Fig 3.**
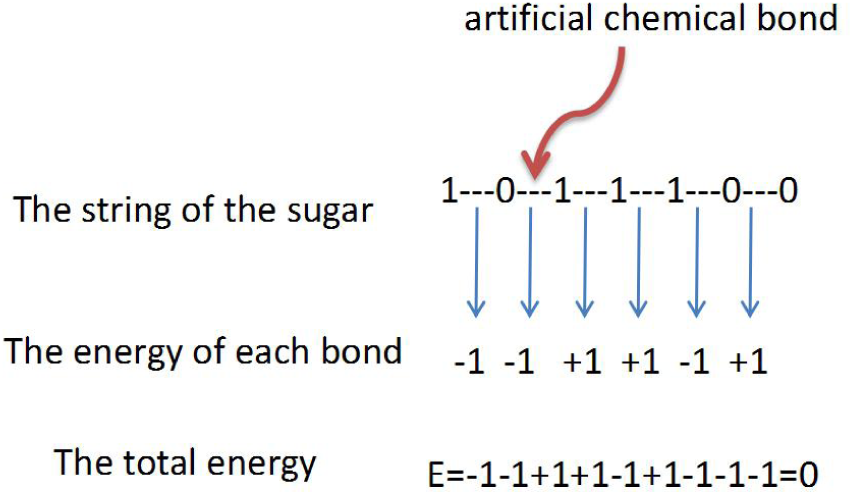
Illustration of artificial potential energy of a string.

In the natural cells, catalytic function of an enzyme takes place only when enzyme, sugar, and ADP collide altogether. To simplify this process, we disentangle them into two independent reactions: first enzyme collides with sugar molecule, and then the sugar molecule with a mark collides with an ADP molecule to transform it into an ATP, as depicted in Fig 4.

**Fig 4.**
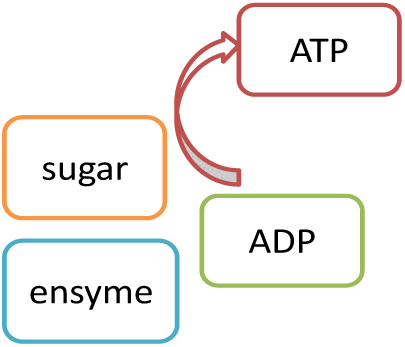
Illustration of the catalytic process.

The energy that a sugar molecule releases when it is catalyzed by an enzyme is determined by the following manner. Each sugar has its own energy which is determined by its string. The enzyme interacts with the sugar molecule and changes its string arrangement, so that the energy of the sugar is changed. If it is decreased by a certain amount, then this reaction happens, and releases the same amount of energy. Otherwise this reaction won’t happen. This process actually mimics the way how a biochemical reaction happens. Detailed description is in the method part.

The dynamics of the artificial biochemical reaction is accomplished as following. We simply use a one dimensional array to store all the molecules. That is a proportion of enzymes, a proportion of sugar molecules, and a proportion of ATP and ADP molecules. We use a scan cycle to update the states of all the molecules just as what we did in the artificial genome to implement the concurrency of the process.

The first state is the existence of the molecule. Enzymes in the array degrade just as TFs on the genome does. Each sugar molecule is refreshed after a fixed number of scan cycles to maintain a sustained nutrient supply. ATPs and ADPs are immortal, just constantly transforming into each other during metabolism. The detailed realization is in the method part.

The second state is the reaction state of the molecule. When an enzyme molecule is scanned, it would randomly select one molecule out of all the molecules in the array. If it is a sugar, then the enzyme will react with the sugar as described before. If not, it will repeat this process till an up limit number is reached. When a sugar molecule is scanned, it will randomly select an ATP molecule in the same manner. Once an ADP molecule is selected, and the sugar has already reacted with an enzyme (we record the number of energies which have been released), the ADP will be transformed into an ATP molecule.

At last, we add another aspect of the ATP molecule. To our knowledge, the ATP molecule is not stable; it has a fixed rate of transforming into ADP molecules. So we also set a fixed decay rate of ATP to ADP. This addition makes the whole system to be dissipative. This means that even if all the transcription activities are halted and consumes no energy, the cell will also dissipate and use all its energy up.

## 4. System integration

With these artificial biochemical reaction rules, the metabolic process is possible of taking place. Without appropriate regulation, however, the reactions serve no biologically meaningful function and most probably are not sustainable. So we need to integrate the metabolic subsystem with the artificial genome subsystem, so that the whole system would realize autonomous metabolic process. Fig. 5 demonstrates the relationship of these two subsystems.

**Fig.5.**
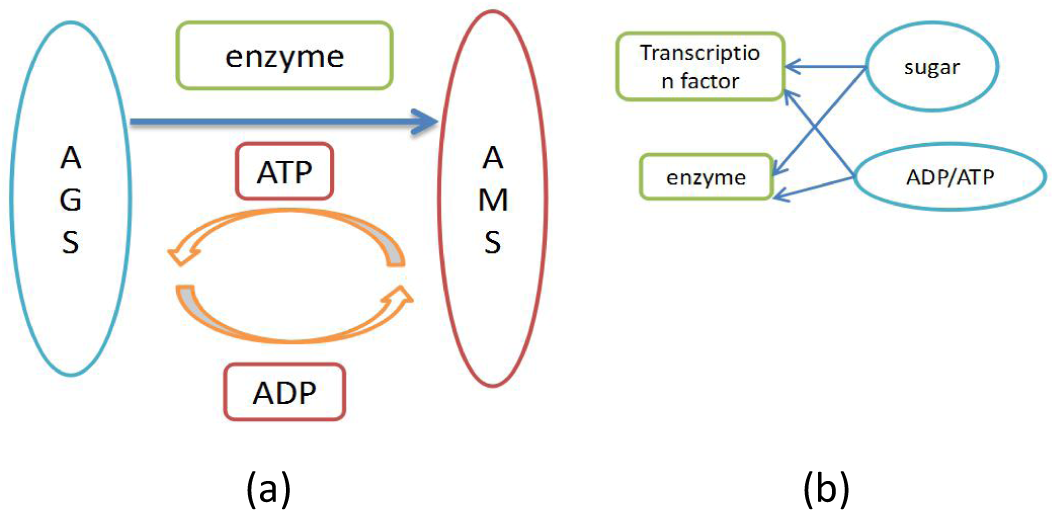
(a)artificial and catabolic dynamical system(b) Illustration of all possible allosterical regulation channel.

The artificial genome subsystem provides enzyme in appropriate times for the metabolic process. All transcription process on the artificial genome consumes energy. For each scan cycle, the transcription of a gene proceeds for one digit if one molecule of ATP is available. We do this by randomly select one molecule in the metabolic molecule array. If an ATP is selected, then it provides one unit of energy needed for the transcription and then transformed into an ADP molecule. If not, the transcription process would halt for on scan cycle. In this way, the time needed for making of a protein is none deterministic. It relays on the availability of ATP molecules.

Transcription factors are also putted into the metabolic array after they are made. But their stay is transient, only a fixed short time, then it would go back to search for its binding position on the DNA. The addition of this process is to make it possible that transcription factors have the ability to take feedback information from the molecules in the catabolic process.

We maintain the entire 4 possible feedback channel waiting for the evolution to decide which one takes place and effect. The negative feedback to the transcription factor is to transform it into a new string, which is irreversible. The positive feedback is just to maintain its existence. The negative feedback of the enzyme is to make it unable to react with sugar. The negative feedback is reversible: the positive feedback would restore its ability of reacting with sugar. Fig.5(b) is the illustration of all the allosterical regulation channel. Take the channel form ATP to enzyme as an example. ATP can either activate the enzyme or deactivate the enzyme, which is determined both by the sequence of ATP and enzyme.

We set an environment where the supply of nutrition is always ample, so that the flux of sugar molecule is sustainable and stable. We accomplish this goal by the following method. The initial life times of sugars are evenly distributed random numbers from 1 to a small up limit. After a sugar’s timer is counted down to 0, it would be cleared and a new sugar molecule would be replenished, with a fixed life time of the up limit.

As we have mentioned, each time an ATP provides energy for the transcription of a protein, it is transformed into an ADP. And here the ADP would be transformed into ATP if a sugar is catalyzed by an enzyme. So the total number of ADP and ATP is constant. They are the energy current of the whole catabolic process. The key of catabolic function lies on the existence of ATP and ADP molecules. We also find that without them, it is hard to form feedback mechanism in catabolic process, which will be discussed in method section.

## 2. The mutational mode of artificial model

As the initial condition of our system is random, the emergence of the artificial metabolic system relies on mutational model of the artificial genome. Ohno [8] proposed the theory that new functions originates from gene duplication in 1970s. Subsequent researchers proposed various kinds of theories based on it. So aside from single nucleotide polymorphism, we also include various kinds of genome rearrangement mode.

**Fig.2.**
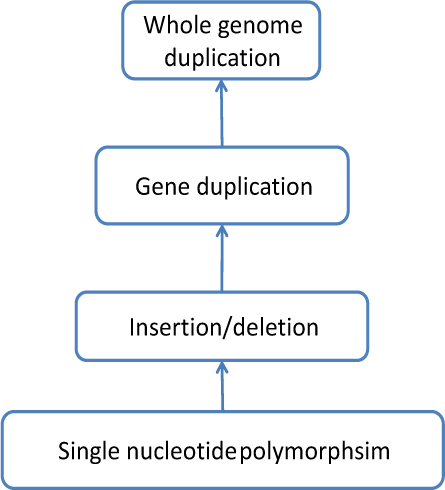
Illustration of hierarchical mutational mode.

We set a hierarchical structure of genome rearrangement, depicted in fig 2. We also include a mechanism of inserting randomly generated genes, mimicking the action of transposons. The mutational rate of a nucleotide is not fixed [10] during evolution. In our model, each nucleotide has its own mutational rate and changes gradually. These settings may give our system the ability to evolve into biological meaningful structures.

## 5. The fitness function

We use evolutionary algorithm to evolve the system. The initial condition of the artificial genome is random, so there is no predetermined gene or regulation relationship. So if this system can evolved into some kind of GRN structure, which regulates some kind of stable catabolic process, then it would shed light on the formation of GRN as well as the metabolic pathway in natural cells.

We chose the life span as fitness value for the evolutionary computation. It is the time during which the system runs continuously. The operation stops when the energy of the system drops to 0 or reaches a maximum value.

We give constant supply of sugar molecule to the system. The transcription process consumes energy while the metabolic process harvests energy. If the system evolve into appropriate structure which can make the consumption and absorption of energy well balanced and its activity sustainable, then the system can maintain long life span.

When one of the following conditions is reached, the system stops its activity.

1: No ATPs exist, which means that energy is depleted.
2: No ADPs exist, which means that there are too much energy storage.
2: All genes are off, which means that the activity of the system is stalled.

## 6. Results of evolutionary algorithm

We set the population to be 60. The initial length of the artificial genomes are all set to be 1000. Here we show one typical result. After 180 generations of evolution, the artificial metabolic system successively emerged. Fig. 8(a) ws the evolution of the life span. In the beginning, the life span lingers around 0 for about 100 generations. This indicates that no biologically meaningful structure has emerged. From about generation 100 to 140, the life span begins to grow. After about 140 generations, the maximum life time soars to about 2 million. This means that some individuals have evolved into very stable metabolic structure. But the average life time is much shorter, about tens of thousands of cycles. This means that the life time among the population has huge gaps. We can also see that after 140 generations, the life time is under intensive fluctuations, which means that the fitness landscape is rather ragged.

**Fig.8.**
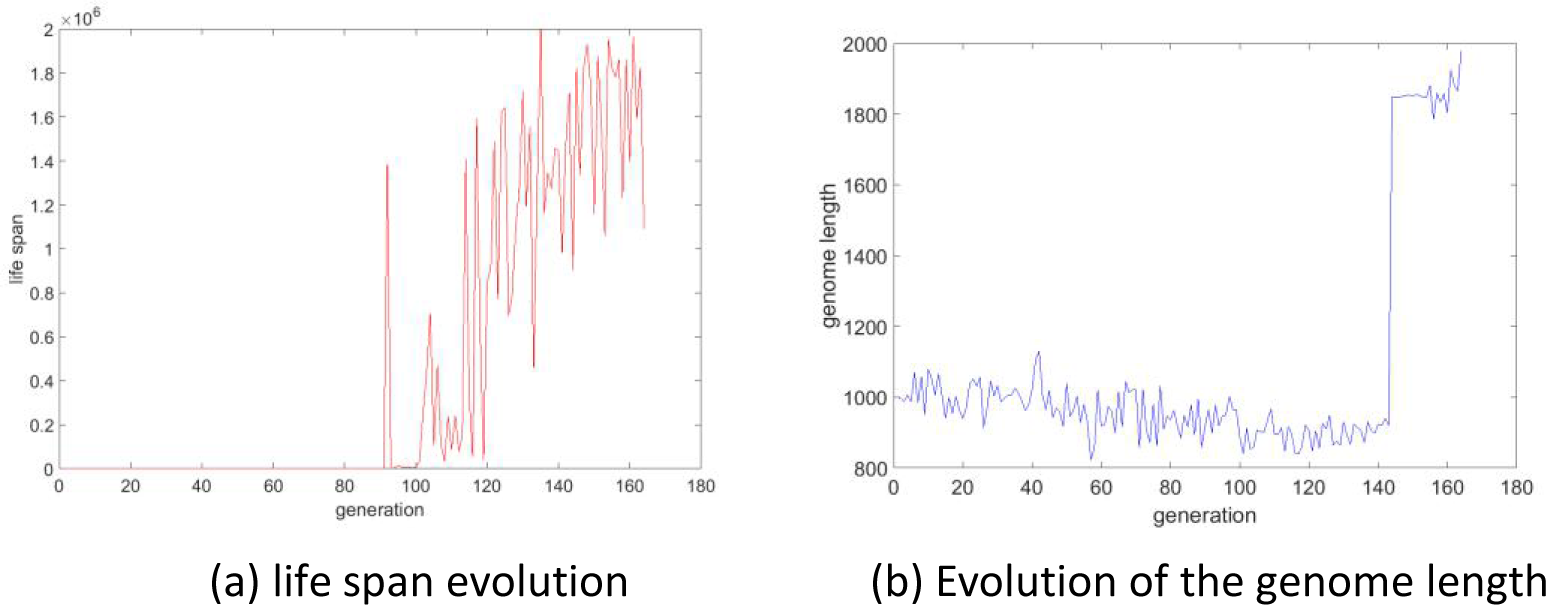
evolution of life span and genome length.

We want to see how our hieratical mutation takes place. We can see in Fig8(b) in the 140^th^ generation, a whole genome duplication event happened. Then the length of the genome fluctuates around about 2000 and is relatively stable. This indicates that whole genome duplication may have influenced the formation of the system. Detailed analysis of this influence will be discussed in the forthcoming paper.

Fig. 10 is the dynamics of one individual after 180 generations. Tracking the feedback manner of the system, we find that ATP molecules all have negative feedback to enzyme molecules, and ADP molecules all have positive feedback to them. This is the ultimate reason of the stable metabolic process. The number of ATPs is dominant, so most of the enzyme molecules are suppressed, but as soon as some ADPs are created caused by energy consumption, the ADPs immediately feedback this information to enzymes and activate them, then leads to the increase of ATP molecules. We can see that in Fig. 10(d), the number of enzyme molecules vibrates around about 25. The number of activated enzymes is around 5, not exceeding 10. This stable vibration is due to the feedback mechanism of ATP and ADP molecules. The whole feedback system thus maintains the stable catabolic process.

**Fig. 10.**
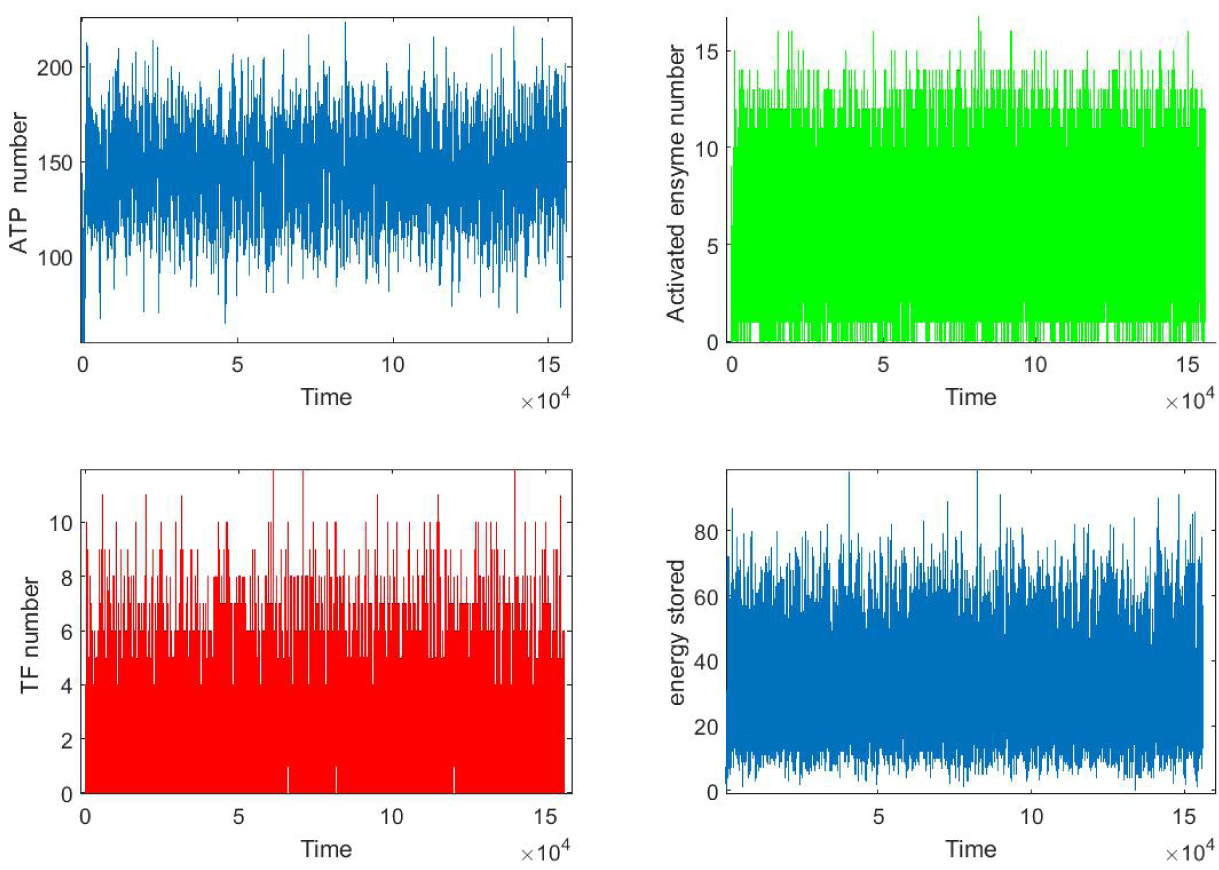
Dynamics of artificial metabolic system. (a) Evolution of the number of ATP molecules. (b) Evolution of activated enzymes. (c) Evolution of the number of TFs (d) Evolution of energy unites stored in sugar molecules.

## 7. Analysis of the artificial metabolic system evolved

Our mean goal is to make the metabolic system emerge from random initial conditions, and see how it is formed during evolution. As we seem to successfully evolve a system of stable catabolic process, we can go back and track down the formation process of the GRN, which is impossible in the real evolution of natural organisms.

Fig 10 displays the TF regulation network from the best fit metabolic system in the 180 generation. Giving the robustness of the system, to our surprise, there are only 7 TF genes in this network. Although there are only 7 genes, their regulation relationship is complex. So we analyze small motifs in this network. We find that there are 3 cycles in this network, and one feed forward motif. These motifs are all well established structures with biological meaningful functions, and spread in all sorts of biochemical pathways. This indicates that the structure computed by our method resembles some characteristics of the real biological system. So inversely, the evolutionary process of our artificial metabolic system may shed light on the formation of the real one in natural cell.

**Fig. 10.**
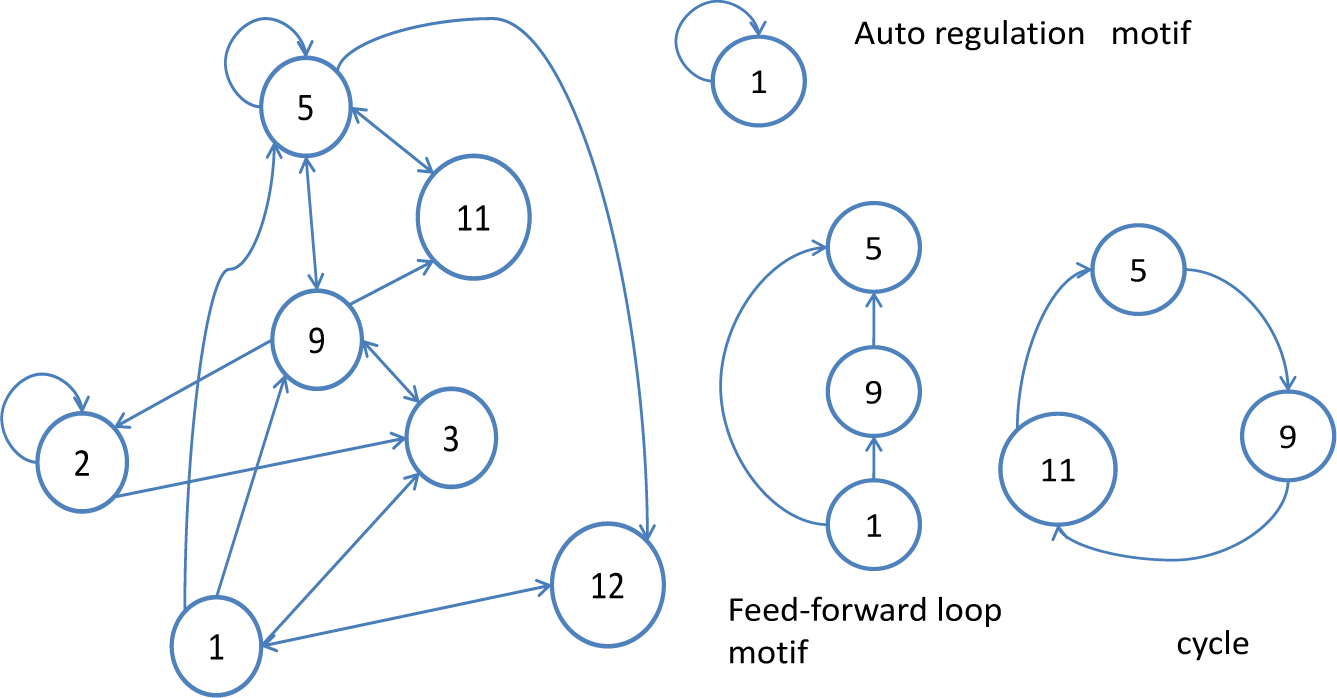
Illustration of the TF regulation network from one metabolic system.

So next,we retrieve the evolutionary history of the network. In the figure below is the GRN network in 25^th^, 50^th^,75^th^ and 100^th^ generation.

## 9. Discussion

From system level perspective, the artificial metabolic system demonstrates some interesting characters. It evolves fast, can evolve to very stable metabolic process, and the regulation network is biological meaningful. We think that the reason may be three folds: First, the hierarchical mutational structure helps the genome sequence to explore the possible GRN space in a very efficient way. Second, as we have mentioned, the incorporation of interacting molecules including enzymes, transcription factors, sugars, ATPs and ADPs. This incorporation makes the functional part of the system possess varies kinds of means to accomplish the metabolic task and in the same time effectively transmit information to the regulatory process. Finally is that these two subsystems run in parallel, and are in constant exchange of material, energy and information. To summarize these 3 factors, we can say that the whole system is autonomous. That is to say, it regulates its own biochemical process.

The role of ATP and ADPs is important, but we can regard them to be one aspect of autonomous system. That is to say, the transmission of energy does not depend on the computer to transform abstract numbers, but solely on basis of the most primitive way of using the transmission of molecules to do this. Although this way seems to be inefficient from the perspective of a computer simulator, form the view of an organism, this way actually generates more possibilities of information transmission and feedback mode. And this phenomenon may occur recurrently in an autonomous system. The artificial genome mode is an example, we bores all the troubles to establish a string to generate genes, rather than directly create genes abstractly using simple nodes and edges. But this bothering is worth it: we obtain abundant ways of form new genes and their regulation relationship, and we harvest very plastic mutational mode. So in the future, we will research in to this direction and analyze how an autonomous factor helps a system to evolve. We hope to get some theoretical results.

Our system can be used as a bench mark for various kinds of theories discussing the evolutionary process of biological processes. Such as the area of how new genes are generated and obtain new functions, what is the role of gene duplication or other form of rearrangement of the genome to the evolution of an organism. Due to space limitation of this article, we just start it a little to make more exciting analysis to come.

We also developed the mathematical model of our system in the method part. We hope that with the help of mathematical models, we can describe biologically system in a more abstract way, and perhaps deduce more profound theoretical results about it.

## Method

### 1.1 artificial genome model

Denote the artificial genome as *G, G*_*i*_ is its ith element. We have *G*_*i*_ ∈{0,1}. Define a substring of *G* as *G*(*i,l*), which means that the substring starts at *G*_*i*_, and has a length of *l*. On the genome, there are a set of genes and its transcription regulation zones. Denote this set as *M, M*_*i*_ is its ith element. *M*_*i*_ = (*z* _*i*_, *g*_*i*_). *z* _*i*_ is the ith transcription zone segment, and *g*_*i*_ is the ith gene segment. We have *z*_*i*_ = *G*(*I*_*z*_ (*i*), *L*_*z*_ (*i*)), *g*_*i*_ = *G*(*I*_*g*_ (*i*), *L*_*g*_ (*i*)). *I*_*z*_ maps the order *i* of the transcription zone to its starting position on the genome and *L*_*z*_ maps *i* to the length of the zone. *I*_*g*_, *L*_*g*_ have the similar function for the ith gene. Define a function Γ, which can map a string on the genome to the set of the substrates of this string. So we have Γ(*z*_*i*_) = {*j* | *G*_*j*_ ∈ *z*_*i*_}. Likewise, Γ(*g*_*i*_) = {*j* | *G*_*j*_ ∈ *g*_*i*_}.

Then we define a function which maps the position on the genome to the substrate of gene or transcription zone it belongs, we have:

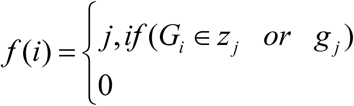

Define a set of all the positions of the transcription zone segment 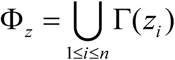, where *n* is the whole number of genes. And likewise, define a set of all the positions of the gene segment 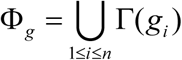. At time t, the segment which is the concatenation of all the TFs which binds to *z*_*i*_ is denoted as *u*_*i*_ (*t*). We use a hash function *H* to map *u*_*i*_ (*t*) to {0,1}. *H* (*u*_*i*_ (*t*)) = 0 means that TFs in this zone deactivates the downstream gene. *H* (*u*_*i*_ (*t*)) = 1 means that TFs in this zone activates the downstream gene. The hash function is selected such that the probability of mapping a string to 0 and 1 is almost equal.

Then we define a random number *F*_*C, K* (*t*)_: Ω → *E*. Ω={0,1},E={0,1}, *E* = 1 means that an ATP happens to available and provides one unit of energy, *E* = 0 means that no ATP is available. *C* is the capacity of the reaction container, *K*(*t*) is the number of ATPs in the container. At each round of scan, the activated gene requires 1 unit of energy from ATP. If it is available, then transcription proceeds for one step, otherwise the transcription is paused. This will be explained further in the following. Now, we have:

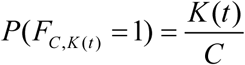

So, this probability varies as *K*(*t*) varies.

### 1.2 realization of concurrency

Define a time line *T, T*_*i*_ is its ith element. We have *T*_*i*_ ∈{*N*}. There is a one to one correspondence between *G*_*i*_ and *T*_*i*_. The state of *T* is updated after each round of scan. We have:

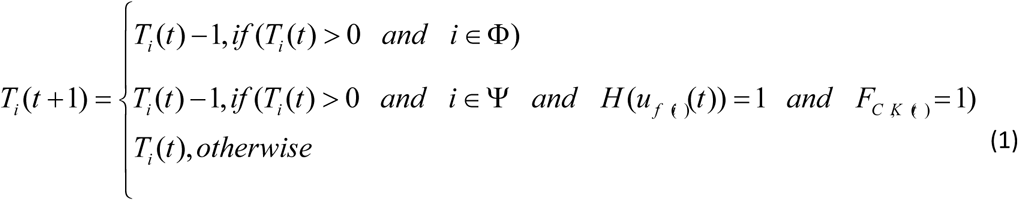

Notice that for each round of scan, *F*_*C, K* (*t*)_ updates only once when the first digit of the *g*_*i*_ is scanned and remain fixed.

The formulation (1) only defined how timers are counted down. There is another formulation which defines how timers are set. The timers are set by events. So next we define how events are triggered.

First we define the event that a new round of transcription is triggered. We define this event as *E*_*trans* _ *start*_ (*i, t*). *i* is the ith position on the artificial genome, *t* is the time. *E*_*trans* _ *start*_ (*i, t*) = 1 means that the a new transcription process is triggered at time t in position *i*. *E*_*trans* _ *start*_ (*i, t*) = 0 means that transcription process is not triggered or an old round of transcription process is not finished yet.

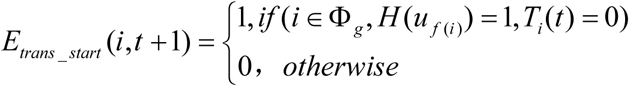

Next, we define the event that a gene has just been thynsersized and bind to a certain place on the genome. We denote this event as *E*_*trans* _ *end*_ (*i, t*). *E*_*trans* _ *end*_ (*i, t*) = 1 means that this event happen, and *E*_*trans* _ *end*_ (*i, t*) = 0 means that this event does not happen. We have:

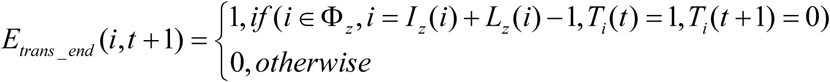

If *E*_*trans* _ *end*_ (*i, t*) = 1, then it immediately triggers the binding of the thynsersized protein to its TF binding site. There may be multiple binding sites available. Next, we define a set of potential binding sites of this TF. First we define a function Ξ, giving a candidate string *S*_*candi*_ it maps all identical strings to this string on the genome, we have:

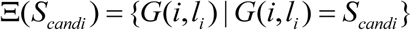

If *G*(*i, l*_*i*_) ∈Ξ{*S*_*candi*_}, then it is possible that the TF *S*_*candi*_ may bind to *G*(*i, l*_*i*_). But it is also possible that *G*(*i, l*_*i*_) does not belong to the transcription zone, or it has already been bound by another TF. So we define a subset of Ξ(*S*_*candi*_) at time *t*, denoted as

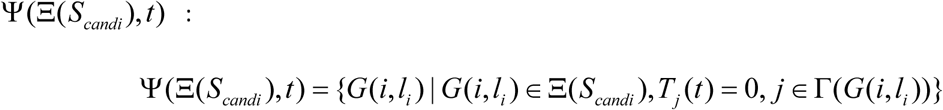

So this set includes all possible binding places of the candidate transcription factor. As we know, the TF can only bind to one place. So we define a random variable 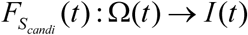,

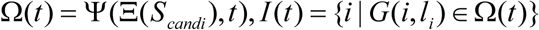

*I*(*t*) is the set of the first positions of all possible binding sites. Here we make the possibility that the TF binds to one of them equally, so we have:

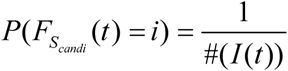

where #(*I* (*t*)) means the number of elements in set *I* (*t*).

So, we have the following laws of event triggered resettings of *T*_*i*_ at time *t*:

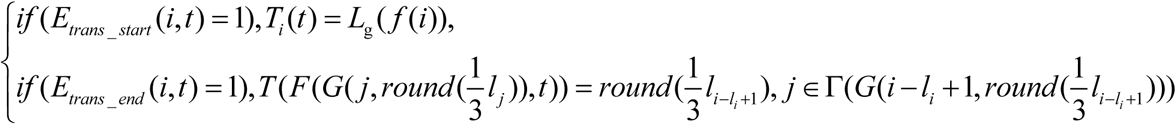

Notice that immediately after *T*_*i*_(*t*) is set to be a positive number, it then will be decreased by following scan cycles. *round* 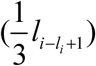 is actually the life span of the TF which has bound to the transcription regulation zone. After this life span is decreased to 0, then this TF is automatically degradated.

#### 2. the metabolic process

Now we describe how the enzyme changes the arrangement of the sugar string.

Suppose that the string length of enzyme and the sugar is the same. Define the sugar sting as *S*. *S*_*i*_ is its ith element. We have *S*_*i*_ ∈{0,1}. The string of enzyme is denoted as *E*, and *E*_*i*_ ∈{0,1} in the same manner. Define an ordered set Γ which records the positions of 1s in *E*. Γ={*j* | *E* _*j*_ = 1} and Γ_*i*_ < Γ _*j*_ if *i* < *j*. Define another ordered set Θ which records the positions of 0s in *E*. Θ = {*j* | *E*_*j*_ = 0} and Θ_*i*_ < Θ _*j*_ if *i* < *j*. Define the third ordered set Π, Π = Γ ∪Θ. Suppose that the length of Γ is *l* and the length of Θ is *r*. We have Π_*i*_ = Γ_*i*_ and Π _*j* +*l*_ = Θ _*j*_. Define a permutation

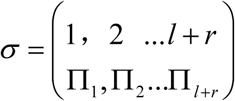

We use this permutation to permutate the string of the sugar molecule. We define a function ψ to compute the energy of a sugar string. So the energy released denoted by Δ is:

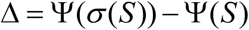

If Δ > 0, then the biochemical reaction happens, otherwise it won’t. It should be mentioned that the transformed sugar string can be transformed again by the same method. This way, our system permit multiple step metabolic process. This allows complex metabolic path way to form.

At last, we should mention that the only predetermined thing is the sequence of the sugar; we set it to be ‘000000111111’. So there is also no predetermined catabolic process, we only determine the interaction rules of the molecules in the array.

## Acknowledgement

This work is supported by Chinese National Natural Science Foundation, Grant number 61702314.

